# Higher habitual FODMAP intake is associated with lower body mass index, lower insulin resistance and higher short-chain fatty acid-producing microbiota in people with prediabetes

**DOI:** 10.1101/2022.10.26.513956

**Authors:** NHS Chu, J He, J Ling, K Leung, RCW Ma, J Lee, J Varney, JCN Chan, JG Muir, E Chow

## Abstract

**Aims/hypothesis:** The quantity and quality of FODMAPs can alter the relative abundance of gut microbiota with metabolic consequences although similar data are lacking in people with prediabetes. We investigated associations between habitual FODMAP contents, gut microbiota and glucose/insulin responses in subjects with prediabetes.

**Methods:** In this prospective cross-sectional study, ninety-eight subjects with impaired glucose tolerance (IGT) (mean age: 57±7 years, 43 % men) had assessment of body composition, 6-point oral glucose tolerance tests (OGTT), Homeostatic Model Assessment of Insulin Resistance (HOMA-IR) and 3-day dietary intake. We analysed faecal samples in a sub-group of 20 subjects with IGT and 10 subjects with normal glucose tolerance by 16S rRNA microbiome analysis.

**Results:** Obese subjects with IGT had the lowest daily FODMAP intake compared with their non-overweight and non-obese counterparts (5.7 (3.9-7.9) vs 7.1 (5.0-11.3) vs 9.9 (4.1-22.4) g/day, p=0.024) despite having similar total daily energy intake. Total content of FODMAPs was negatively correlated with body fat. After adjustment for age and gender, total FODMAPs were negatively associated with BMI and HOMA-IR. This remained significant after adjustment for macronutrients and physical activity (p=0.032 and p=0.036 respectively). FODMAP contents were strongly associated with short-chain fatty acid (SCFA)-producing bacteria, such as *Lactobacillus* (p=0.011), *Akkermansia muciniphila* (p=0.012), and *Bifidobacterium longum* (p=0.010), the abundance of which were negatively correlated with 2-hr plasma glucose (r = -0.524, p =0.003).

**Conclusion:** In individuals with IGT, higher habitual FODMAP intake was associated with lower body fat and insulin resistance and increased abundance of SCFA-producing bacteria, calling for interventional studies to evaluate the effects of FODMAP intake in prediabetes.

## Introduction

Prediabetes is a high-risk state with 5-10% of affected people progressing to diabetes annually. The prevalence of prediabetes is increasing worldwide with more than 470 million people estimated to have prediabetes by 2030(1, 2). Dietary intervention is effective in prevention of diabetes and changes in gut microbiota may be implicated (3, 4). Lifestyle, dietary and genetic factors can influence fat accumulation. While low-grade inflammation in visceral adipose tissue can increase insulin resistance(5, 6), intestinal microbiome can modulate lipid metabolism through multiple mechanisms including changes in short-chain fatty acid (SCFA) (7, 8).

FODMAPs (fermentable oligosaccharides, disaccharides, monosaccharides, and polyols) are a group of fermentable short chain carbohydrates that are metabolised by gut microbiota. A low FODMAP diet can reduce the symptoms of irritable bowel syndrome by 50-80%(9, 10). However, this is accompanied by increased abundance of *Bacteroides* and reduced abundance of *Bifidobacterium* (11, 12) and *Akkermansia muciniphila* known to have favourable metabolic consequences. Through fermentation, these microbiota generate active metabolites which can alter the balance of deconjugated secondary bile acids(13), SCFA(14), lipopolysaccharide (LPS)(7) and incretin secretion(15, 16), which are implicated in glucose and lipid metabolism.

Supplementation with individual FODMAP items as prebiotics have been shown to alter body composition, gut hormones, and gut microbiota in animal studies. Fructo-oligosaccharides added to a high fat diet reduced body fat mass and adiposity in rats(17). Galactooligosaccharides (GOS) supplementation lowered LDL-cholesterol, increased *Bifidobacterium* and reduced *Clostridium* in mice fed with high fat diet(18). In another rodent study, GOS increased the incretin hormones, glucagon-like peptide-1 (GLP-1) and peptide YY (PYY) and abundance of health-promoting *Bifidobacterium*(19). In a double-blind, parallel randomized clinical trial (RCT) involving 44 overweight/obese subjects with prediabetes, 15g of GOS daily when added to regular diet increased *Bifidobacterium* by 5-times but did not alter peripheral insulin sensitivity, SCFA, LPS or inflammatory markers(20). However, compared with prebiotic supplementation, FODMAPs from whole foods might exert additional effects on whole gut transit time and motility(21) and interact with other gut components resulting in different effects.

In a study involving subjects with impaired glucose tolerance (IGT) participating in a diabetes prevention program, we documented their habitual FODMAP content at baseline. In a subgroup of 20 subjects, we also examined their microbiome. In this analysis, we examined the cross-sectional associations between habitual FODMAP content, body mass index (BMI), body composition, insulin resistance and their associations with gut microbiome before intervention. We hypothesised that higher habitual FODMAP consumption is associated with greater abundance of SCFA-producing bacteria. Further, individual FODMAPs may affect gut microbiota profiles and orchestrate different changes in metabolic phenotypes (22, 23).

## METHOD

### Participants

Participants underwent screening to identify individuals with IGT for participation in a 12-month RCT which evaluates the effects of continuous glucose monitoring (CGM) as an adjunct to lifestyle modification for prevention of glycemic deterioration (NCT NCT04588896). In this investigator-initiated, single-centre study conducted at the Prince of Wales Hospital (PWH), the teaching hospital of the Chinese University of Hong Kong, participants were identified from attendees at the PWH medical outpatient clinics or self-referral through advertisements. Inclusion criteria included age range of 18-65 years, body mass index (BMI) range of 18-40 kg/m^2^, non-pregnant or lactating state, no history of diabetes and treatment with glucose lowering or weight-reducing drugs. Subjects who participated in a weight-reducing program within 3 months of screening were also excluded. All participants underwent a 75 g oral glucose tolerance test (OGTT) after an overnight fast. Glycaemic status and type 2 diabetes were defined according to the American Diabetes Association (ADA) criteria (2009): 1) Normal glucose tolerance (NGT): fasting plasma glucose (FPG) <5.6 mmol/L and 2-hour PG<7.8 mmol/L; 2) impaired fasting glycemia (IFG): 5.6 mmol/L≤FPG≤6.9 mmol/L; 3) impaired glucose tolerance (IGT): 7.8 mmol/L≤2-hour PG<11.1 mmol/L; 4) diabetes: FPG≥7 mmol/L or 2-h PG≥11.1 mmol/L. The study was approved by the CUHK Clinical Research Ethics Committee.

### Anthropometrics, body composition, and physical activity

Anthropometric measures were taken with the subjects wearing light clothing and without shoes. Body weight and body composition (body fat percent) were assessed by the bioelectric impedance analysis system (Tanita; Model: TBF-410 Body composition analyser)(24, 25). Waist and hip circumference (cm) were measured using a standard, retractable, non-metallic tape measure placed round the waist at the level of the umbilicus and across the largest part of the buttocks, and below the iliac crest, respectively. Height was measured with a stadiometer to the nearest 0.1 cm for calculation of BMI. According to the World Health Organization (WHO) Asian classification for obesity(26), the cut-off point for overweight was 23 kg/m^2^ and 27 kg/m^2^ for obesity. Therefore, we defined normal weight as 18-22.9 kg/m^2^, overweight as 23-26.9 kg/m^2^, and obesity as ≥27 kg/m^2^. Physical and activity levels were recorded using the International Physical Activity Questionnaires (IPAQ) (Chinese version)(27).

### Biochemical profiles

A 75-g oral glucose tolerance test (OGTT) was performed in the morning after an overnight fast. Blood samples were collected through a venous catheter from an antecubital vein into vacutainer tubes containing EDTA (ethylenediamine tetraacetic acid) at 0, 15, 30, 60, 90, and 120 minutes for the determination of plasma glucose and at 0, 30 and 120 minutes. Plasma CP concentration was measured by radioimmunoassay (Novo Nordisk, Copenhagen, Denmark). The lowest detection limit was 0.1 nmol/L with an intra-assay coefficient of variation (CV) of 3.4%, and an inter-assay CV of 9.6%(28).

We computed the steady state of insulin resistance and dynamic indices of beta cell function using fasting CP and PG values. Insulin resistance (HOMA2-IR) and insulin secretion (HOMA2-2B) were calculated using The Homeostasis Model Assessment (HOMA2) Calculator v2.2.3. downloaded from http://www.dtu.ox.ac.uk(29). We analysed the HOMA2-IR score as continuous value with high value indicating increased insulin resistance. We calculated PG and CP area under the curve (AUC) during OGTT. Fasting blood samples were collected for measurement of HbA1c, triglycerides, total cholesterol, high-density lipoproteins (HDL-C) and low-density lipoprotein (LDL-C) cholesterol.

### Dietary evaluation

Eligible and participants who gave written consent were instructed to document their habitual dietary intake prospectively using food records over three days before randomization. Each of the food records constituted one weekend and two weekdays to accurately capture variations in food intake between weekends and weekdays. A 3-day food record has been shown to adequately rank the habitual intake of FODMAPs by an individual (30) in a non-interventional setting. Subjects were asked to provide details including quantities of all consumed meals and beverages according to standardized portion sizes published by the Centre for Food Safety Hong Kong(31). Upon return of the food records, the research dietitian carefully assessed the records to ensure no missing food item with confirmation by another research nutritionist. The food records were analyzed for energy, macronutrient, and total dietary fiber content using a nutritional analysis programme (eSHA Food Analysis and Labelling Software and FoodWorks 10 – Xyris). The contents of individual FODMAPs, excess fructose (i.e. fructose in excess of glucose), lactose, fructans, GOS and polyols (sorbitol and mannitol), were calculated using the published Monash University FODMAP composition database (The Monash FODMAP calculator)(32).

### Gut microbiota analysis

Fecal samples were collected from a random subset of 30 subjects including 20 IGT participants and 10 control subjects with NGT. Fecal samples were collected at home in sterile plastic tubes and stored at -4°C in a provided ice-containing flask before being returned to the Centre within 2 hours which was then stored at −80°C for batch analysis. The microbial DNA was extracted using the QIAamp PowerFecal Pro DNA Kit and the V3–V4 region (300bp*2) of the 16S rRNA gene was amplified and sequenced on a HiSeq Illumina. All association analyses were performed using normalized absolute abundancies, i.e. counts that were normalized by cumulative sum scaling (CSS) in R using the metagenomics Seq package. We described the percentage abundances and relative abundances of the bacterial genera in these study subjects. The Shannon diversity index was calculated the proportional abundance of species using the R package *vegan*.

### Statistical analysis

#### Descriptive analyses

For comparisons, Student’s t-test, Mann-Whitney U test, Chi-square (χ^2^), Fisher’s exact test, or Analysis of variance (ANOVA) were used as appropriate. Post-hoc Tukey tests were performed for between-group comparisons if the overall results were significant. The non-parametric Kruskal-Wallis test was used to compare between-group differences in dietary intake of macronutrients and individual FODMAPs. If significance was detected, post-hoc analysis of significant values was performed using the least significant difference test between the two subject groups. Values were reported as mean (95% confidence interval) for parametric data or median [interquartile range] for non-parametric data.

### Association analyses

In the whole cohort of 98 subjects with IGT, we used Spearman correlation analysis to study relationships between total and individual FODMAP intake with body composition (BMI and total body fat) as well as that between total and individual FODMAP item. We then constructed multivariate linear model of the FODMAP item versus body composition indices with age and sex as covariates (model 1). In model 2, we additionally included daily consumption of macronutrients and total sugar intake as covariates and in model 3, we included physical activity. We also examined association of FODMAP intake with HOMA-IR adjusted for aforementioned covariates. In the subset of individuals with gut microbiota (n=30) analyses, we performed correlations analyses between FODMAP and microbiota at phylum, genus, and species level.

Statistical analysis of all data was performed using Statistical Package for Social Sciences version 26.0 (SPSS Inc., Chicago, IL, USA). The R software package was used to calculate the correlation of FODMAPs with microbiota and insulin indexes.

## Results

### Baseline characteristics of the IGT cohort

Between December 2020 and 2021, we screened 303 subjects for eligibility using 75 g OGTT. Of these, 100 subjects were confirmed to have IGT (two IGT subjects were excluded in the analysis) and 152 had normal 2-hour PG and 43 had undiagnosed diabetes. Seven subjects withdraw consent and one subject was not fulfilled the criteria (Supplementary 1). The number of cases during the study period determined the sample size (33, 34) and we excluded the missing data on food dairies of the subject. (Supplementary 1). All IGT subjects along with 10 subjects with NGT were instructed to complete a 3-day food records and other measurements. The compliance of 3-day food records including two consecutive weekdays and one weekend was 99%.

Table 1 summarises the characteristics of the study population. Amongst 98 IGT participants included in this analysis, 80% were either obese or overweight, 43% were men with a mean±SD age of 57±7 years, and BMI of 26±4.6 kg/m^2^. The median of total FODMAPs in the IGT cohort was 6.8 (IQR: 4.5-10.9) g/day and the distribution of individual FODMAPs are shown in Figure 1. Amongst individual FODMAP items, fructans had the highest consumption followed by lactose and excess fructose. The median intake of GOS was 0.37 g/day.

**Table 1.** Baseline demographics in subjects with IGT analysed by BMI categories.

| *Variable* | *Total Population (n=98)* | *Normal (n=16)* | *Overweight (n=37)* | *Obesity (n=45)* | *P-value* | *Missing data (%)* |
| --- | --- | --- | --- | --- | --- | --- |
| *Age, years* | 57±7 | 58±7.0 | 59±6 | 55±8 | 0.063 | 0 |
| *Male, n (%)* | 42 (43%) | 6 (38%) | 18 (51%) | 17 (38%) | NA | 0 |
| *Weight, kg* | 68±13 | 57±5 | 68±7 | 78±12 | <0.0001 | 0 |
| *Height, cm* | 163±7.7 | 163±6 | 164±7 | 160±9 | 0.196 | 0 |
| *Waist circumferences, cm* | 91±11 | 80±4 | 92±5 | 101±8 | <0.0001 | 0 |
| *Hip circumferences, cm* | 99±8 | 91±4 | 98±3 | 106±6 | <0.0001 | 0 |
| *BMI, kg/m<sup>2</sup>* | 26±4.6 | 21±1.1 | 25±1.1 | 30±2.8 | <0.0001 | 0 |
| *Systolic blood pressure, mmHg* | 133±16 | 128±16 | 134±16 | 138±17 | 0.059 | 0 |
| *Diastolic blood pressure, mmHg* | 84±10 | 80±8 | 86±10 | 85±12 | 0.110 | 0 |
| *Body fat, %* | 32.2±9.9 | 24.1±6.7 | 28.9±6.3 | 37.4±8.8 | <0.0001 | 1 |
| *Fasting plasma glucose, mmol/L* | 5.4±0.6 | 5.3±0.5 | 5.4±0.5 | 5.4±0.5 | 0.815 | 0 |
| *1hr plasma glucose, mmol/L* | 11.0±1.7 | 10.8±1.6 | 10.9±1.8 | 11.1±1.6 | 0.821 | 1 |
| *2hr plasma glucose, mmol/L* | 8.6±1.4 | 8.4±1.2 | 8.8±1.3 | 9.0±1.1 | 0.285 | 0 |
| *Fasting plasma C-peptide, pmol/L* | 600±294 | 392±155 | 602±221 | 774±276 | <0.0001 | 2 |
| *2hr plasma C-peptide, pmol/L* | 3037±913 | 2572±757 | 3134±1056 | 3477±943 | 0.006 | 2 |
| *HOMA-IR2(cp)* | 1.36±0.68 | 0.87±0.33 | 1.36±0.51 | 1.74±0.63 | <0.0001 | 2 |
| *HOMA- $\beta$ (cp)* | 99.0±35.2 | 78.5±31.3 | 99.6±26.1 | 120.5±34.5 | <0.0001 | 2 |
| *C-peptide AUC, pmol/L* | 230592±71812 | 177787±52562 | 224170±71768 | 255049±67476 | 0.001 | 3 |
| *Physical Activities* |  |  |  |  |  |  |
| *Vigorous, min/week* | 0(0-53) | 0(0-75) | 0(0-98) | 0(0-11) | 0.367 | 3 |
| *Moderate, min/week* | 0(0-120) | 60(0-180) | 40(0-180) | 0(0-40) | 0.027 | 4 |
| *Light, min/week* | 180(90-360) | 195(41-585) | 210(99-420) | 163(65-240) | 0.257 | 11 |
| *Sedentary, min/day* | 300(210-480) | 300(180-360) | 285(180-435) | 360(240-540) | 0.233 | 17 |

Mean±SD, median (interquartile range)

**Figure 1.**
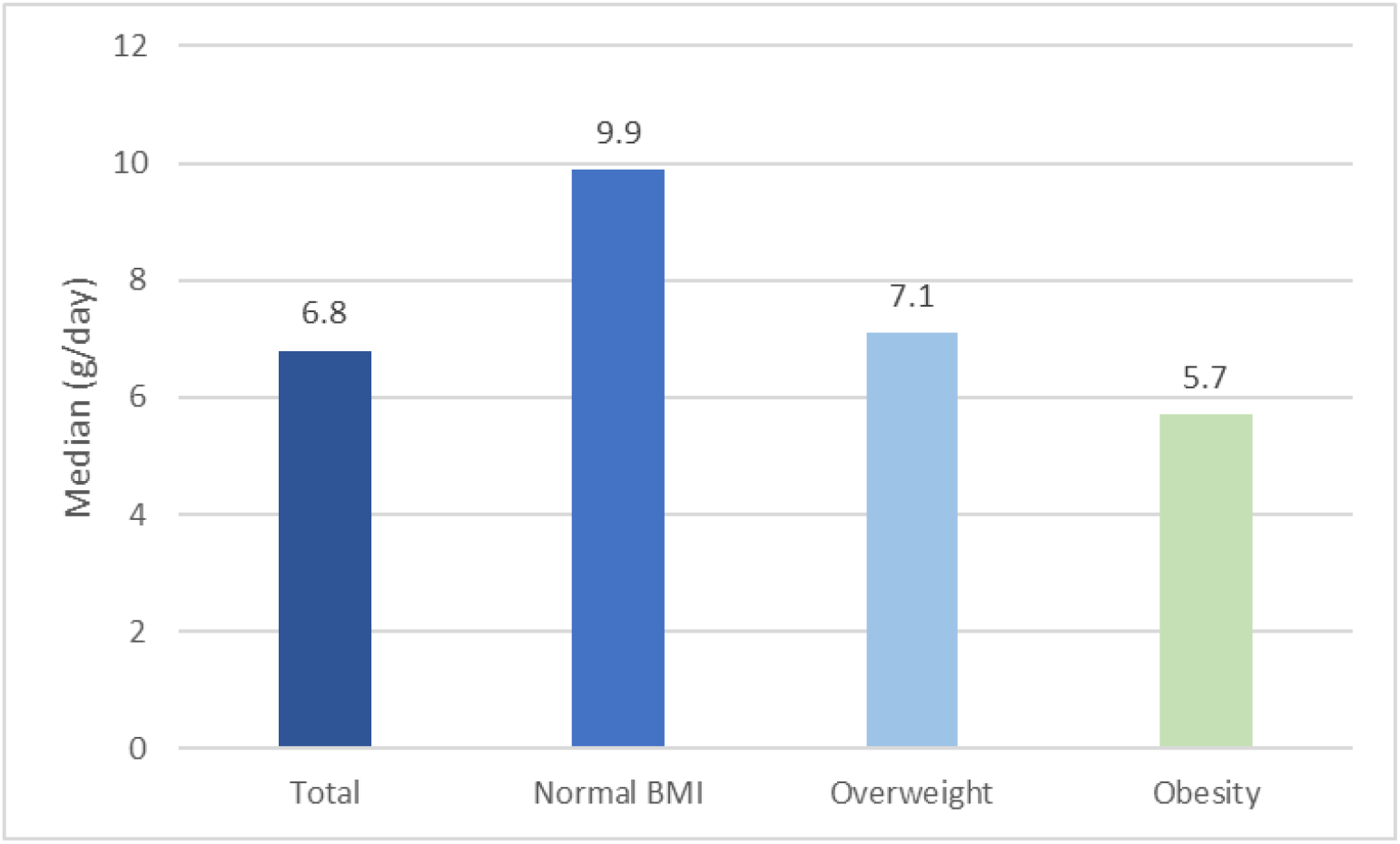
The median of individual FODMAP contents in the IGT study population. Total study population, normal body mass index (BMI), overweight and obesity groups. (From left to right).

### FODMAP and body composition

There were no between-group differences in macronutrients intake except for a lower fiber intake amongst obese individuals (Table 2). Physical activity levels were similar amongst all 3 groups. Daily habitual FODMAP intake decreased progressively with increasing BMI, ranging from 9.9 (4.1-22.4) g/d in normal-BMI group, 7.1 (5.0-11.3) g/d in overweight-group to 5.7 (3.9-7.9) g/d in obese subjects (p=0.024 Kruskal Wallis Test). For individual FODMAPs; GOS (0.22 vs 0.48 g/day, p=0.08) and mannitol intake was lower (0.09 vs 0.29g/day, p=0.048) in the obese than the normal-BMI group. There were no significant differences in excess fructose or lactose intake analysed BMI categories. (Table 2)

**Table 2.** Median consumption of macronutrients and FODMAPs by BMI categories.

| | Normal (n=16)<br>BMI >18 and <23<br>kg/m<sup>2</sup> | Overweight(n=37)<br>BMI 23-26.9 kg/m<sup>2</sup> | Obesity(n=45)<br>BMI $\geq 27$ kg/m<sup>2</sup> | P value |
| --- | --- | --- | --- | --- |
| Energy, kcal/day | 1938(1561-2396) | 2153(1816-2419) | 1931(1614-2143) | 0.167 |
| Protein, g/day | 97(70-103) | 90(81-111) | 87(70-104) | 0.321 |
| Fat, g/day | 81(70-96) | 87(63-96) | 82(63-96) | 0.385 |
| Carbohydrates, g/day | 214(151-251) | 229(195-267) | 208(170-261) | 0.374 |
| Sugar, g/day | 42(32-56) | 46(36-60) | 45(28-66) | 0.611 |
| Fiber, g/day | 15.6(8-19) | 13.5(10-15) | 8.7(6-12) | <0.0001 |
| Total FODMAPs, g/day | 9.9(4.1-22.4) | 7.1(5.0-11.3) | 5.7(3.9-7.9) | 0.024 |
| Excess Fructose<sup>#</sup>, g/day | 1.1(0.4-2.3) | 1.2(0.4-1.6) | 0.5(0.3-1.3) | 0.160 |
| Polyols*, g/day | 1.0 (0.2-1.8) | 0.9 (0.4-1.6) | 0.2 (0.1-0.8) | 0.003 |
| Fructans, g/day | 2.3(1.5-3.6) | 2.2(1.6-2.7) | 1.7(1.2-2.4) | 0.015 |
| GOS, g/day | 0.48(0.27-1.31) | 0.61(0.30-1.20) | 0.22(0.13-0.43) | <0.0001 |
| Lactose, g/day | 3.2(0.4-11.0) | 1.4(0.5-4.4) | 1.8(0.2-4.0) | 0.235 |

*#Excess fructose is defined as fructose minus glucose, \*Polyols is the sum of mannitol and sorbitol*
*No missing data on nutritional data.*

In these 98 subjects with IGT, FODMAPs was negatively correlated with total body fat (r = −0.324, p=0.001). For individual component of FODMAPs, GOS was negatively correlated with BMI (r = −0.379, p<0.0001), waist (r = −0.267, p=0.008), and hip circumference (r = −0.334, p=0.001). Mannitol was also negatively correlated with BMI (r = −0.294, p =0.003), waist (r = −0.233, p=0.021), and hip circumference (r = −0.300, p=0.003). Lactose was negatively correlated with body fat (r = −0.282, p=0.005) (Table 4). There were no significant correlations for other individual FODMAPs such as excess fructose, sorbitol or fructans. In multivariate analysis, there was negative association between total FODMAP intake and BMI (β = −0.172, p = 0.008) following adjustment for age and gender. These negative associations persisted following adjustment for macronutrients (β = −0.192, p = 0.004) and physical activity (β = −0.197, p = 0.004) (Table 3).

**Table 3.** Multivariate analysis of association between total FODMAP consumption and BMI, body fat and insulin resistance.

| Dependent variable | Beta coefficient | 95% CI | P value | Adjusted R<sup>2</sup> |
| --- | --- | --- | --- | --- |
| **BMI (kg/m<sup>2</sup>)** |  |  |  |  |
| Base model | -0.173 | [-0.300 to -0.047] | 0.008 | 0.063 |
| Model 1 | -0.196 | [-0.328 to -0.064] | 0.004 | 0.059 |
| Model 2 | -0.196 | [-0.326 to -0.065] | 0.004 | 0.130 |
| **% Body fat** |  |  |  |  |
| Base model | -0.272 | [-0.503 to -0.042] | 0.021 | 0.516 |
| Model 1 | -0.253 | [-0.500 to -0.007] | 0.044 | 0.510 |
| Model 2 | -0.306 | [-0.562 to -0.051] | 0.019 | 0.530 |
| **HOMA IR** |  |  |  |  |
| Base model | -0.024 | [-0.044 to -0.004] | 0.020 | 0.070 |
| Model 1 | -0.023 | [-0.044 to -0.002] | 0.032 | 0.084 |
| Model 2 | -0.024 | [-0.047 to -0.002] | 0.036 | 0.102 |

BMI, body fat and HOMA-IR were included as dependent variable with daily FODMAP as independent variable

**Table 4.** Spearmen correlation coefficients of individual and total FODMAPs and anthropometrics, biochemical profiles and insulin secretion/resistance indices.

| Correlation coefficient (Spearman) | Fructans | GOS | Excess Fructose | Lactose | Sorbitol | Mannitol | Total FODMAPs |
| --- | --- | --- | --- | --- | --- | --- | --- |
| Age, years | r=0.114<br>p=0.263 | r=0.150<br>p=0.141 | r=0.015<br>p= 0.883 | r= -0.024<br>p=0.815 | r= -0.096<br>p=0.347 | r=0.022<br>p=0.827 | r=0.020<br>p=0.846 |
| Sex | r= -0.024<br>p=0.814 | r=0.087<br>p=0.392 | r= -0.043<br>p=0.674 | r= -0.271<br>p=**0.007** | r= -0.059<br>p=0.566 | r=0.136<br>p=0.181 | r= -0.225<br>p=**0.026** |
| Weight, kg | r=-0.074<br>p=0.468 | r= -0.292<br>p=**0.004** | r= -0.019<br>p=0.854 | r= 0.100<br>p=0.329 | r= -0.017<br>p=0.871 | r= -0.318<br>p=**0.001** | r= -0.001<br>p=0.991 |
| Waist circumference, cm | r= -0.145<br>p=0.154 | r= -0.267<br>p=**0.008** | r= -0.146<br>p=0.153 | r= -0.073<br>p=0.476 | r= -0.182<br>p=0.072 | r= -0.233<br>p=**0.021** | r= -0.213<br>p=**0.035** |
| Hip circumference, cm | r= -0.103<br>p=0.315 | r= -0.334<br>p=**0.001** | r= -0.129<br>p=0.207 | r= -0.047<br>p=0.644 | r= -0.103<br>p=0.313 | r= -0.300<br>p=**0.003** | r= -0.164<br>p=0.107 |
| BMI, kg/m<sup>2</sup> | r= -0.170<br>p=0.094 | r= -0.379<br>p<**0.0001** | r= -0.147<br>p=0.148 | r= -0.041<br>p=0.688 | r= -0.194<br>p=0.055 | r= -0.294<br>p=**0.003** | r= -0.193<br>p=0.057 |
| Body fat, % | r= -0.144<br>p=0.159 | r= -0.167<br>p=0.102 | r= -0.099<br>p=0.334 | r= -0.282<br>p=**0.005** | r= -0.112<br>p=0.276 | r= -0.037<br>p=0.717 | r= -0.324<br>p=**0.001** |
| Fasting glucose, mmol/L | r= 0.017<br>p=0.869 | r= 0.044<br>p=0.666 | r= -0.016<br>p=0.878 | r=0.197<br>p=0.052 | r= 0.018<br>p=0.863 | r= -0.016<br>p=0.874 | r=0.151<br>p=0.139 |
| Fasting C-peptide, pmol/L | r= -0.166<br>p=0.107 | r= -0.287<br>p=**0.005** | r= -0.023<br>p=0.826 | r= -0.035<br>p=0.733 | r= -0.160<br>p=0.119 | r= -0.178<br>p=0.083 | r= -0.133<br>p=0.195 |
| HOMA-IR | r= -0.163<br>p=0.113 | r= -0.282<br>p=**0.005** | r= -0.014<br>p=0.890 | r= -0.026<br>p=0.802 | r=0.150<br>p=0.145 | r= -0.175<br>p=0.087 | r= -0.123<br>p=0.232 |
| HOMA-beta | r= -0.180<br>p=0.080 | r= -0.283<br>p=**0.005** | r= -0.039<br>p=0.707 | r= -0.132<br>p=0.201 | r= -0.185<br>p=0.070 | r= -0.165<br>p=0.109 | r= -0.221<br>p=**0.030** |
| 1hr PG, mmol/L | r= -0.083<br>p=0.418 | r= -0.062<br>p=0.545 | r= -0.099<br>p=0.333 | r= 0.166<br>p=0.104 | r= -0.022<br>p=0.834 | r= -0.030<br>p=0.771 | r= 0.039<br>p=0.704 |
| 2hr PG, mmol/L | r= 0.029<br>p=0.780 | r= 0.015<br>p=0.880 | r= -0.084<br>p=0.409 | r= -0.015<br>p=0.881 | r=0.000<br>p=0.998 | r= 0.027<br>p=0.788 | r= -0.065<br>p=0.524 |
| 2hr plasma C-peptide, pmol/L | r= -0.226<br>p=**0.027** | r= -0.192<br>p=0.061 | r= 0.054<br>p=0.603 | r= -0.044<br>p=0.671 | r= -0.109<br>p=0.290 | r= -0.150<br>p=0.146 | r= -0.127<br>p=0.216 |
| Plasma CP AUC, pmol/L | r= -0.218<br>p=**0.032** | r= -0.209<br>p=**0.041** | r= 0.031<br>p=0.763 | r= -0.046<br>p=0.655 | r= -0.135<br>p=0.189 | r= -0.173<br>p=0.092 | r= -0.139<br>p=0.178 |
| Total Cholesterol, mmol/L | r= 0.111<br>p=0.275 | r= -0.005<br>p=0.961 | r= -0.056<br>p=0.581 | r= 0.150<br>p=0.140 | r=0.018<br>p=0.859 | r=0.104<br>p=0.310 | r=0.114<br>p=0.265 |
| HDL-C, mmol/L | r= 0.128<br>p=0.210 | r= 0.192<br>p=0.059 | r= 0.132<br>p=0.196 | r= -0.070<br>p=0.491 | r=0.083<br>p=0.419 | r=0.222<br>p=**0.028** | r=0.066<br>p=0.521 |
| LDL-C, mmol/L | r= 0.116<br>p=0.261 | r= -0.001<br>p=0.990 | r= -0.047<br>p=0.649 | r= 0.172<br>p=0.093 | r=0.014<br>p=0.891 | r=0.082<br>p=0.425 | r=0.137<br>p=0.182 |
| Triglycerides, mmol/L | r= -0.022<br>p=0.828 | r= -0.144<br>p=0.157 | r= -0.089<br>p=0.381 | r= -0.016<br>p=0.872 | r= -0.134<br>p=0.190 | r= -0.050<br>p=0.628 | r= -0.098<br>p=0.338 |

PG=plasma glucose; CP=C peptide; HOMA=Homeostasis model assessment for insulin resistance (HOMA-IR) and beta cell function (HOMA-beta) derived from fasting PG and CP

### Association between FODMAP and indices of insulin secretion and resistance

There was no correlation between 2-h PG and FODMAP contents (Table 4). For individual FODMAPs, GOS was negatively correlated with HOMA-IR and HOMA-β (r = −0.282, p=0.005 and r = −0.283, p=0.005). No signification correlations were observed between FODMAPs and early or late C peptide response during OGTT. In multivariate analysis, total FODMAPs were significantly associated with HOMA-IR and HOMA-beta (β = −0.024, p=0.020 and β = −1.446, p=0.010) after adjustment of age and gender. These remained significant following adjustment for macronutrients (β = −0.023, p=0.032 and β = −1.457, p=0.013) and physical activity (β = −0.024, p=0.037 and β = −1.688, p=0.011).

### Associations between dietary FODMAPs and microbiota

Gut microbiota profiles were available for a subset of participants (10 = NGT) and (20 = IGT). The IGT and NGT groups had similar age and BMI (Supplementary 2). Both groups had similar intake of macronutrients with the IGT group having lower habitual FODMAP content including GOS, sorbitol and mannitol than the NGT group (p<0.05) (Supplementary 3). Compared to subjects with NGT, the IGT group had a higher relative abundance of phylum *Desulfobacterota* (including genus *Bilophila*) and lower relative abundance of *Bifidobacterium longum* at the species level.

FODMAP contents were strongly associated with SCFA-producing bacteria. GOS was positively correlated with *Lactobacillus* (p=0.011) and *Akkermansia* (p=0.012). Fructans was negatively correlated with *Parasutterella* (p=0.042). Mannitol was negatively correlated with *Streptococcus* (p=0.043). Sorbitol was positively correlated with *Sutterella (p=0*.*004)*. Lactose was positively correlated with *Bifidobacterium longum* (p=0.010) and negatively with *Escherichia/Shigella (p=0*.*034)*. There was no correlation between excess fructose and target microbiota (Figure 2).

**Figure 2.**
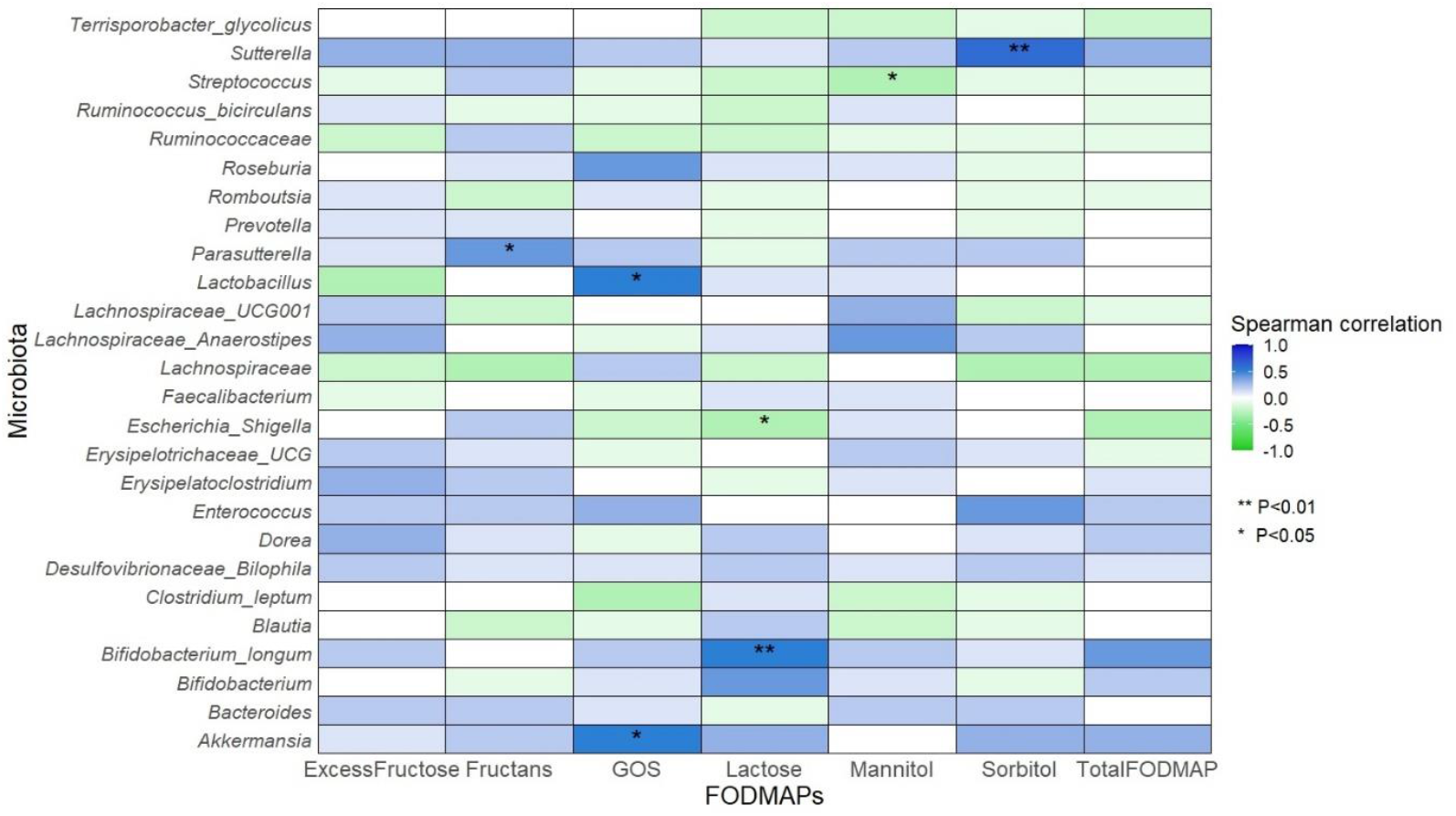
Spearman correlation of individual FODMAPs and microbiota. ** P<0.01, *P<0.05. Blue colour shows positive correlation and green colour shows negative correlation.

HOMA-IR was positively correlated with *Bilophila* (p=0.022). Both HOMA-IR and HOMA-beta were negatively correlated with *Lachnospiraceae* (p=0.001 and p=0.023 respectively). Two-hour plasma C-peptide was negatively correlated with *Bifidobacterium longum* (p=0.016), *Lactobacillus* (p=0.020) and *Lachnospiraceae* (p=0.012). The AUC of plasma C-peptide was negatively correlated with *Bifidobacterium longum* (p=0.029) and *Lactobacillus* (p=0.011), but positively correlated with *Bilophila* (p=0.039). (Supplementary 4)

### Interaction between gut microbiota

Apart from absolute differences, we also observe changes in relative abundance of different microbiota. *Bifidobacterium longum* was associated with lower abundance of *Bilophila* (p=0.003) and increased abundance of *Anaerostipes* (p=0.004) and *Clostridium leptum* (p=0.008). Besides, *Lactobacillus* was associated with increased abundance of the family *Lachnospiraceae* (p=0.006) and reduced abundance of the family *Ruminococcaceae* (p=0.038) and the genus *Escherichia/ Shigella* (p=0.040).

## Discussion

In this prospective study including 98 Chinese individuals with IGT, cross-sectional analysis indicated independent association of high FODMAP consumption with lower BMI, body fat and insulin resistance after adjustment for macronutrients and fibre intake and physical activity, with the strongest associations observed with GOS. In a subgroup of 20 subjects with IGT along with 10 subjects with NGT, FODMAP contents were strongly associated with SCFA-producing bacteria, such as *Lactobacillus, Akkermansia muciniphila* and *Bifidobacterium longum*. These observational data support our hypothesis that habitual FODMAP content may modulate adiposity, insulin resistance, possibly mediated through changes in gut microbiota.

We observed a low consumption of FODMAP in our IGT subjects (median 6.8 (4.5-10.9) g/day). Although not directly comparable, patients with IBS from the United Kingdom, Australia, United States, Sweden, and New Zealand reported a daily intake of 16-31g (35-39). We and others have shown that South and East Asians had generally lower FODMAP consumption than their European counterparts in both normal population and those with IBS (40, 41). Overall, our Chinese subjects with IGT had low consumption of milk and dairy products (26 ml/day of milk) and high intake of rice (217 g/day). The latter accounted for 57.1% of carbohydrate intake. They also consumed fruits and vegetables with low FODMAP content, such as oranges (consumed by 47.6% of study population), Choy sum (56%) and lettuce (34%). Despite these differences, there are no reports comparing habitual FODMAP between Asians and European population with prediabetes or diabetes.

Amongst Chinese subjects with IGT, the obese group had the lowest intake of total FODMAP content (obese: 5.7 g/day; overweight: 7.1g/day and non-obese: 9.9 g/day). Interestingly, we found robust associations between habitual consumption of FODMAPs and BMI, body fat and insulin resistance, independent of total calorie or macronutrient intake and physical activity. In a subgroup analysis of 20 subjects with IGT and 10 subjects with NGT, we also observed close associations between gut microbiome and body composition. Increased consumption of FODMAPs was associated with absolute increase and relative abundance of SCFA-producing microbiota (42-44). Here, SCFA can modulate the activity of transcription factor peroxisome proliferator-activated receptor-γ (PPAR-γ). The latter regulates adipocyte differentiation with reduced ectopic fat accumulation and improved lipid and glucose metabolism (45, 46). Secondly, these bacterial metabolites, such *as Akkermansia muciniphila* phospholipids which can activate the sympathetic nervous system and restore the activity of gastrointestinal endocrine cells with increased secretion of gut hormones such as PYY, GLP-1 and Cholecystokinin (CKK). These changes in metabolic and hormonal milieu can systemically regulate gluconeogenesis and glycogenolysis in liver and lipolysis from adipose tissues(16, 47, 48), as well as these hormones work on the brain-gut axis to regulate food intake via increasing epigastric fullness and satiety(49, 50).

In support of the increased abundance of health-promoting microbiota, we also found independent associations of individual and total FODMAPs with insulin response and 2-hour PG during OGTT. Besides, daily FODMAP intake was associated with higher abundance of *Bifidobacterium longum*, the latter being associated with lower 2-hr PG level (r = -0.524, p =0.003). Other researchers had demonstrated that microbiome-derived signals could influence adipose tissue biology via suppression of nucleotide-binding oligomerization domain-containing protein 1 (NOD1) with increased functional NOD2 in adipose tissue, resulting in reduced insulin resistance and inflammation(15). In healthy subjects, a 24-hr high FODMAP diet reduced LPS binding protein compared to consumption of a low FODMAP diet. To this end, chronic subclinical inflammation and LPS are known to increase insulin resistance (51).

In our study, GOS was negatively correlated with BMI and insulin resistance and positively correlated with *Akkermansia muciniphila*. There was also reciprocal suppression of harmful bacteria such as *Bilophila* and *Escherichia/ Shigella*, which might contribute to the lower 2-hr PG response associated with high GOS intake. In animal (52, 53) and human trials (54, 55), *Akkermansia muciniphila* was found to be inversely correlated with body fat mass. In a randomized, parallel, double-blind study, consumption of a GOS-containing diet increased the abundance of *Bifidobacterium* sequences and decreased the abundance of *Bilophila wadsworthia* in patients with functional gastrointestinal disorders given a low FODMAP diet for relief of symptoms(56). In our subjects with IGT, the higher abundance of *Bilophila* was positively associated with HOMA-IR (r = 0.424, p = 0.022), fasting PG (r = 0.405, p=0.026) and 2-hr PG (r =0.379, p=0.039). Apart from being a microbiota, GOS is also a major prebiotic which can increase the abundance of *Bifidobacterium* to further reduce metabolic endotoxemia and improve glucose tolerance in human(57). GOS can be a natural extract from indigestible carbohydrate materials or synthetically produced in various forms presented as liquid (syrup), capsule or natural foods. However, GOS as a prebiotic supplementation, had not been shown to improve metabolic indices in human study(20). In our study, we found favourable association of high FODMPA intake including GOS with glucose and insulin metabolism. These differences might be due to slower regional transit time of GOS from natural foods (58, 59) with increased contact time with other gut microbiota and possible interactions with other FODMAPs in contrast to administration of GOS as a prebiotic supplement.

This is the first study investigating the metabolic and microbiota associations of habitual FODMAP in people with prediabetes. Low habitual FODMAP contents may unfavourably influence the diversity of gut microbiome and predispose to metabolic diseases. In this study, we obtained detailed metabolic phenotyping with multiple-point OGTT and documentation of FODMAPs and macronutrients using a 3-day food diary and physical activity using structured questionnaire. However, our study also had limitations. We did not assess body composition using dual energy X-ray absorptiometry (DEXA) although bioimpedance analysis (BIA) is a validated measure of body composition including body fat percentage which closely correlates with that measured by DEXA and Magnetic resonance imaging (MRI)(24, 25). Physical activity was only assessed using self-reported questionnaire while use of sealed pedometers and accelerometers may allow more accurate quantification. Our sample size is relatively small and only a subset had faecal microbiota analysis. Nevertheless, our findings were robust and remained significant following adjustment of macronutrient and physical activity. Since these individuals were recruited for a lifestyle modification program, they might be more motivated with higher health literacy and their diets may not be generalizable. That said, these data were collected at baseline before any intervention. Our results might be confounded by unmeasured variables, such as genetic factors, indigested proteins, glycemic index or load. High FODMAP foods generally have a lower glycemic index with total amount of carbohydrate in a food showing better correlations with diabetes risk than glycemic index or load (60). In our study, total carbohydrate intake was similar amongst the 3 groups analysed by obesity. We also excluded subjects who had recently used antibiotics and probiotics in this cross-sectional analysis. Finally, as association does not infer causation, our findings are hypothesis-generating and future interventional studies are needed to confirm the impact of FODMAP intake on metabolic outcomes in prediabetes.

## Conclusion

In Chinese subjects with IGT, dietary FODMAP intake was low especially in those with obesity. Higher FODMAP intake was associated with lower BMI and insulin resistance as well as increased SCFA-producing bacteria, independent of total energy and macronutrient intake and physical activity. Our findings have potential implications on the use of FODMAP as a specific dietary strategy to prevent or manage diabetes, beyond calorie restriction and low-carbohydrate restricted diets. Our results also support novel mechanisms of diet-microbiota-host interactions underlying cardiometabolic diseases (61). Future studies are needed to better understand how dietary FODMAP components can influence metabolic response and gut microbiome.

## Abbreviations

BIA: bioimpedance analysis
CGM: continuous glucose monitoring
CKK: Cholecystokinin
CP: C-peptide
CSS: cumulative sum scaling
DEXA: Dual energy X-ray absorptiometry
FODMAP: fermentable oligosaccharides, disaccharides, monosaccharides and polyols
GLP1: glucagon-like peptide-1
GOS: Galactooligosaccharides
IGT: Impaired glucose tolerance
LPS: Lipopolysaccharide
NGT: Normal glucose tolerance
NOD: nucleotide-binding oligomerization domain-containing protein
PPYR γ: Peroxisome proliferator-activated receptor-γ
PYY: Peptide YY
rRNA: Ribosomal ribonucleic acid
SCFAs: Short-chain fatty acids

## CONTRIBUTIONS

NHSC and EC conceived the idea of the study, JH, KL, NHSC were involved in data collection and supported by RCWM and JCNC. JL, JL, JV, JGM, NHSC analysed the data. NHSC wrote the first draft of the manuscript. All authors critically reviewed the manuscript and approved the final version.

