## Supplementary for "Higher habitual FODMAP intake is associated with lower body mass index, lower insulin resistance and higher short-chain fatty acid-producing microbiota in people with prediabetes"

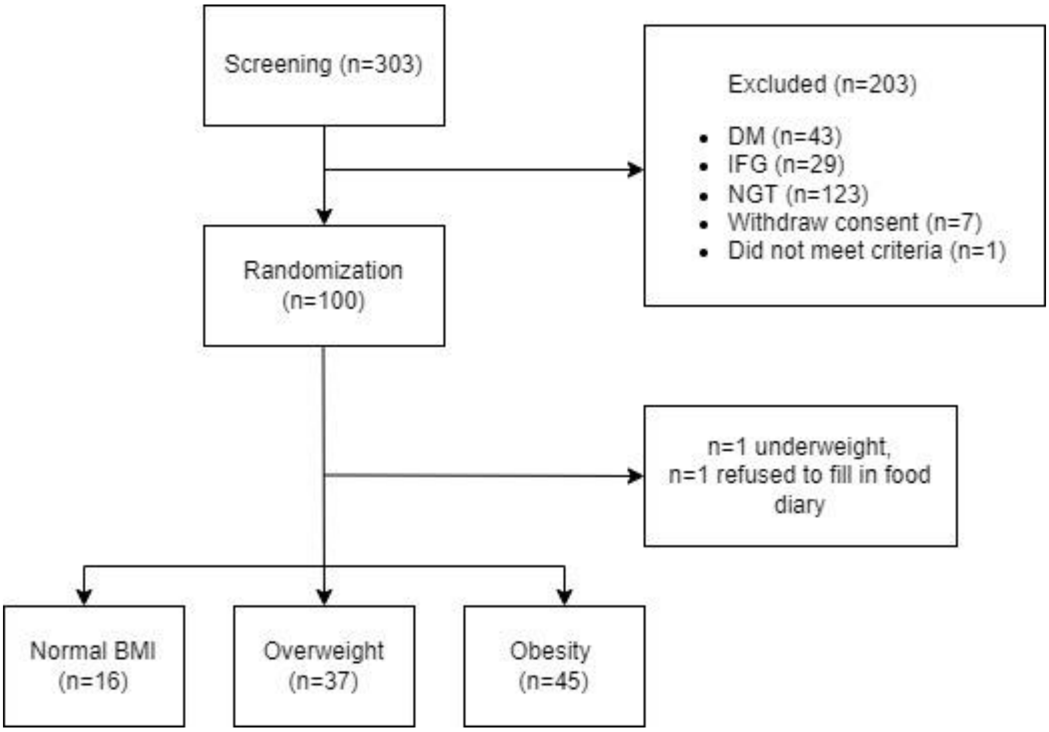


Supplementary 1**.** Flow Chart of the study

Supplementary 2. Baseline demographic data between NGT and IGT. Abbreviations: BMI: body mass index, SBP: systolic blood pressure, DBP: diastolic blood pressure.

| | *Normal glucose tolerance (NGT) n=10* | *Impaired glucose tolerance (IGT) n=20* | *P value* |
|---|---|---|---|
| *Age (years)* | 54.5±9.6 | 59.3±4.6 | 0.166 |
| *Men, N(%)* | 5 (50%) | 10 (50%) | NA |
| *Height (cm)* | 160.0±6.0 | 162.5±6.6 | 0.399 |
| *Weight (kg)* | 58.9±10.9 | 65.7±9.9 | 0.095 |
| *Waist (cm)* | 81.3±8.6 | 89.4±8.1 | 0.017 |
| *Hip (cm)* | 91.4±4.2 | 96.8±6.4 | 0.023 |
| *BMI (kg/m$^2$)* | 22.7±2.5 | 24.8±3.5 | 0.101 |
| *Body fat (%)* | 22.9±5.2 | 29.6±9.1 | 0.040 |
| *Trunk fat (%)* | 26.5±10.3 | 35.8±7.3 | 0.012 |
| *Visceral fat (%)* | 7.6±3.2 | 11.8±4.9 | 0.034 |
| *Fasting PG(mmol/L)* | 4.9±0.5 | 5.6±0.6 | 0.003 |
| *2hr PG (mmol/L)* | 5.7±1.0 | 9.0±0.8 | <0.0001 |
| *Fasting C-peptide (pmol/l)* | 305 (589-224) | 484 (742-315) | 0.003 |
| *2hr C-peptide (pmol/l)* | 1848 (2296-1374) | 3039 (3866-2429) | <0.0001 |
| *HOMA IR* | 0.67 (0.47-1.30) | 1.11 (1.63-0.71) | 0.031 |
| *HOMA B* | 87.5 (66.1-110.9) | 78.2 (57.8-111.4) | 0.769 |
| *C-peptide AUC (pmol/L* | 3103±939 | 4151±1133 | 0.019 |

Supplementary 3. Dietary intake from 3-day food diary records between NGT and IGT. Abbreviations: GOS: galato- oligosaccharides, FODMAP: fermentable oligosaccharides, disaccharides, monosaccharides, and polyols.

| | *Normal glucose tolerance (NGT) N=10* | *Impaired glucose tolerance (IGT) N=20* | *P-value* |
|---|---|---|---|
| *Total energy intake (kcal/d)* | 2143±643 | 1846±384 | 0.122 |
| *Protein (g/d)* | 96±25 | 85±17 | 0.157 |
| *Fat (g/d)* | 89±25 | 75±20 | 0.114 |
| *Carbohydrates (g/d)* | 236±91 | 212±57 | 0.376 |
| *-sugars (g/d)* | 40±26 | 45±21 | 0.556 |
| *Fibre (g/d)* | 15±8 | 14±6 | 0.692 |
| *Total FODMAPs (g/d)* | 8.2 (6.0-10.7) | 10.3 (5.0-13.2) | 0.588 |
| *Excess fructose#(g/d)* | 1.0 (0.4-2.0) | 0.8 (0.5-1.5) | 0.779 |
| *Polyols*(g/d)* | 0.66 (0.07-2.11) | 0.76 (0.43-1.52) | 0.746 |
| *Fructans(g/d)* | 2.1 (1.4-2.9) | 2.1 (1.5-2.9) | 0.983 |
| *GOS(g/d)* | 0.23 (0.11-0.57) | 0.39 (0.25-1.12) | 0.183 |
| *Lactose(g/d)* | 4.0 (2.4-6.4) | 4.0 (0.1-7.4) | 0.713 |

***#Excess fructose is defined as fructose minus glucose, *Polyols is the sum of mannitol and sorbitol***

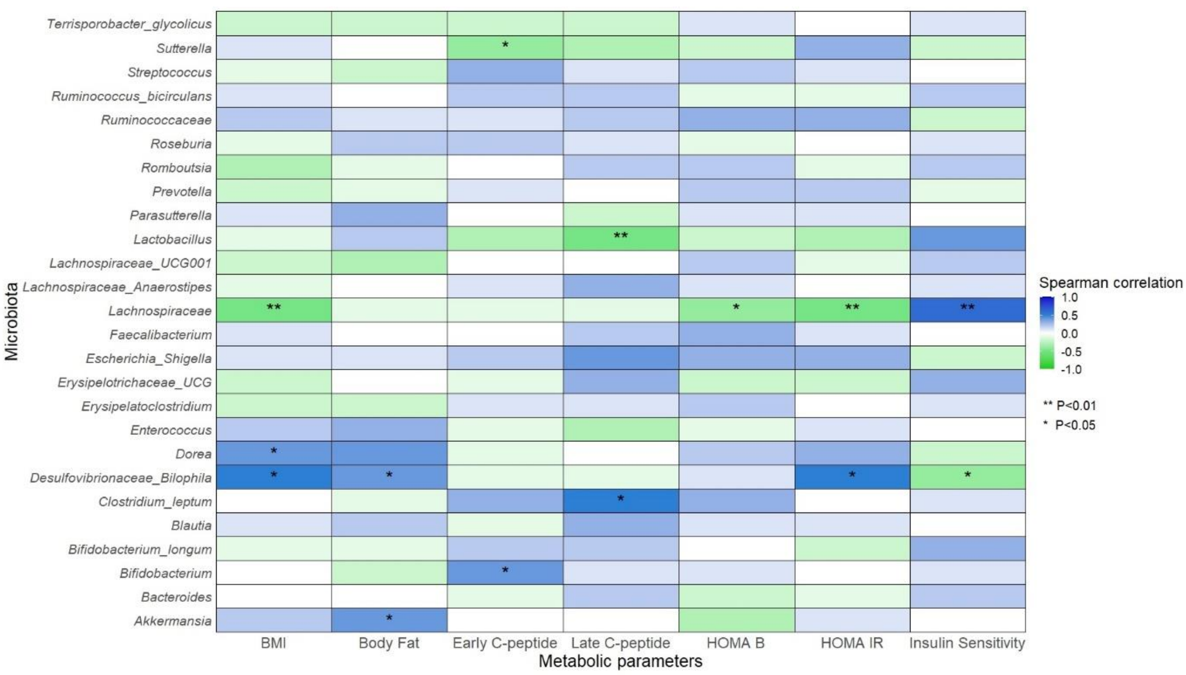


Supplementary 4. Correlation of Homeostatic Model Assessment (HOMA) and gut microbiota
